# Future climate change impacts on anchoveta (*Engraulis ringens*) in the Northern Peru Current Ecosystem

**DOI:** 10.1101/2023.02.14.528548

**Authors:** Ricardo Oliveros-Ramos, Yunne-Jai Shin

**Author notes:** Corresponding author: Ricardo Oliveros-Ramos.

## Abstract

The Northern Peru Current Ecosystem (NPCE) is the most productive ecosystem in terms of fish biomass and sustains the world’s largest small pelagic fishery, the Peruvian anchovy fishery. A cooling of this system has been observed during recent decades but the potential regional impacts of rising atmospheric CO_2_ concentrations on upwelling dynamics and productivity are still unknown. We used the ecosystem model OSMOSE to forecast the impacts of several scenarios of climate change on Peruvian anchovy in the NPCE. The OSMOSE model was forced by plankton production and climate drivers from the Earth System Models IPSL CM5A-LR and GFDL-ESM2M for the period 2009-2100. For each earth system model, representative concentration pathways (RCP) scenarios 2.6, 4.5, 6.0 and 8.5 were run. Our results showed that an optimistic trajectory for anchovy is a reduction of biomass at a rate of 14% per decade until mid-21st century, followed by a collapse and late recovery by the end of the 21st century, while no changes in the spatial distribution of the population were observed. The pessimistic trajectory for anchovy is a reduction of biomass of 22% per decade, with a collapse after 2020 and near extinction by 2060, with a spatial displacement of the population to the south and to more coastal areas. Further research is needed to include additional key environmental variables such as oxygen as well as more realistic fisheries management intervention scenarios to evaluate future options for the sustainability of anchovy resource and fisheries.

## Introduction

The ocean provides about 17% of global animal protein consumed by humans (FAO, 2016) and is a source of livelihoods for millions of people (Sumaila et al., 2012). Coastal marine ecosystems are among the most valuable and heavily used natural systems (Halpern et al., 2008), with 40% of the world’s population within 100 km distance to the coast. The magnitude of projected climate change, in combination with fisheries exploitation, indicates that marine resources will undergo increased pressure in the future (Pörtner et al., 2014; UN, 2016). The assessment of potential future impacts of climate change is required to anticipate the ensuing consequences on ecosystem resilience (Bernhardt and Leslie, 2013) and food security (Barange et al., 2014), being particularly critical in coastal areas which concentrate multiple drivers of change (Bernhardt and Leslie, 2013).

The Northern Peru Current Ecosystem (NPCE) is the most productive coastal ecosystem in terms of fish production and sustains the world’s largest small pelagic fishery, the anchoveta (*Engraulis ringens*) or Peruvian anchovy fishery. This fish production is sustained by the wind-driven upwelling of cold and nutrient replete deep waters near the coast, but this system is also subject to one of the highest environmental variability in the world (Chavez et al. 2008). The anchoveta fishery is highly variable at various time scales and its fluctuations have major societal and economic impacts. A cooling of this system has been observed during recent decades but the potential regional impacts of climate change on upwelling dynamics and productivity are still unknown. The responses of ecosystems to perturbations are difficult to predict because they result from complex interactions among the ecosystem components. This causes enormous challenges for science in terms of prediction capabilities, and for fisheries management in terms of adaptation capacities. In the long term, uncertainties lie in the future state of fish resources which are currently heavily exploited in Peru, like Peruvian anchovy, in the context of climate change and natural environmental variability. Moreover, Earth system models (ESM) used to forecast the impacts of climate change do not provide high resolution outputs that are required to produce reliable forecasts for coastal areas. This provides an additional difficulty to build regional marine ecosystem models as tools to provide scenario-driven projections of future fish dynamics and fisheries production (e.g. Blanchard et al., 2012; Lehodey et al., 2015; Mullon et al., 2016) under climate change.

The objective of the present work is to develop a set of scenarios for Peruvian anchovy under climate change, by using a regional marine ecosystem and multispecies model OSMOSE (Shin and Cury 2001, 2004) for the NPCE (Oliveros-Ramos 2014) which would allow to address direct impacts of climate change on anchovy as well as indirect effects through trophic interactions with the other modelled species of the system. OSMOSE will be forced with two ESMs: IPSL-CM5A (Dufresne et al. 2013, Mignot et al. 2013) and GFDL-EM2M (Dunne et al. 2012, Dunne et al. 2013) and the range of available emission scenarios (four Representative Concentration pathways, RCPs) in order to address the uncertainty due to the climate change scenarios and models used, and to explore the range of variability of possible future responses of anchovy.

## Methods

In order to explore the impacts of climate change scenarios on Peruvian anchovy (*Engraulis ringens*), we used the ecosystem model OSMOSE to reproduce the dynamics of the main marine fish in the Northern Peru Ecosystem including anchovy and, the scenarios of change in climate drivers from the IPSL-CM5A (Dufresne et. al 2013, Mignot et al. 2013) and GFDL-ESM2M (Dunne et al. 2012, 2013) Earth system models. The forecast was done using a 1-way coupling between the physical-biogeochemical model and the upper trophic level multispecies fish model as described in Travers et. al (2009). The period used for the forecast of climate change scenarios was 2006-2100, while the base model was developed for the period 1992-2008 and for the spatial domain extending from 93ºW-70ºW and 20ºS-5ºN (Oliveros–Ramos 2014, see Figure 1). For the hindcast period, the forcing of the OSMOSE model were the plankton fields (phyto and zooplankton, taken from the regional physical and biogeochemical ROMS-PISCES model) and the spatial distribution of fish (estimated using remote sensing environmental inputs and Ecological Niche Models, Oliveros-Ramos 2014). In addition, larval and fishing mortalities were estimated within the model for 1992-2008 using multiple sources of data time series (Oliveros-Ramos et al. 2017) and an evolutionary algorithm (Oliveros-Ramos 2016). For the forecast period (2006-2100), plankton fields were derived from a statistical downscaling and bias correction procedure applied to the outputs from ESMs (Oliveros-Ramos et al. submitted); while the spatial distribution maps of fish were constructed using the same models for the hindcast period but predicted over the future environmental variables. However, larval and fishing mortalities were not available for the future period, and while the latter could be part of a scenario (as a control and policy-intervention variable), we needed to build a statistical model between the historical larval mortalities and the environmental variables available so we could forecast this important input for the model. A conceptual diagram of the modelling approach is shown in Figure 2 and the summary of the models used in Table 1, and the methodological details are provided in the next sections.

**Table 1:** Description of the models used for the construction of climate change scenarios for Peruvian anchovy.

| Model | Description | Reference |
| --- | --- | --- |
| Hindcast (1992-2008) |  |  |
| ROMS-PISCES | ROMS models the 3D physical dynamics of the ocean. The biogeochemical model PISCES simulates the biological marine productivity and describes the biogeochemical cycles of carbon and main nutrients in the ocean. Both models are coupled and produce the dynamics of the low trophic level (phyto- and zooplankton). | Aumont and Bopp, 2006<br>Espinoza et al. 2016 |
| OSMOSE | The higher trophic level model OSMOSE (Object-oriented Simulator of Marine ecOSystems Exploitation) is a spatial multispecies and individual-based model which focuses on fish species. | Shin and Cury (2001, 2004),<br>Oliveros-Ramos (2014) |
| Climate niche model | Statistical species distribution models based on the climate niche ecological theory. | Oliveros-Ramos (2014) |
| Forecast (2009-2100) |  |  |
| OSMOSE | Coupled to the ocean and the plankton dynamics of the IPSL-CM5A and GFDL-ESM2M models. | This study. |
| IPSL-CM5A | ESM developed by the Climate Modeling Center of the Institut Pierre Simon Laplace (France), forecast up to 2099. | Dufresne et. al 2013<br>Mignot et al. 2013 |
| GFDL-ESM2M | ESM developed by the NOAA's Geophysical Fluid Dynamics Laboratory (USA), forecast up to 2099. | Dunne et al. 2012<br>Dunne et al. 2013 |

**Figure 1.**
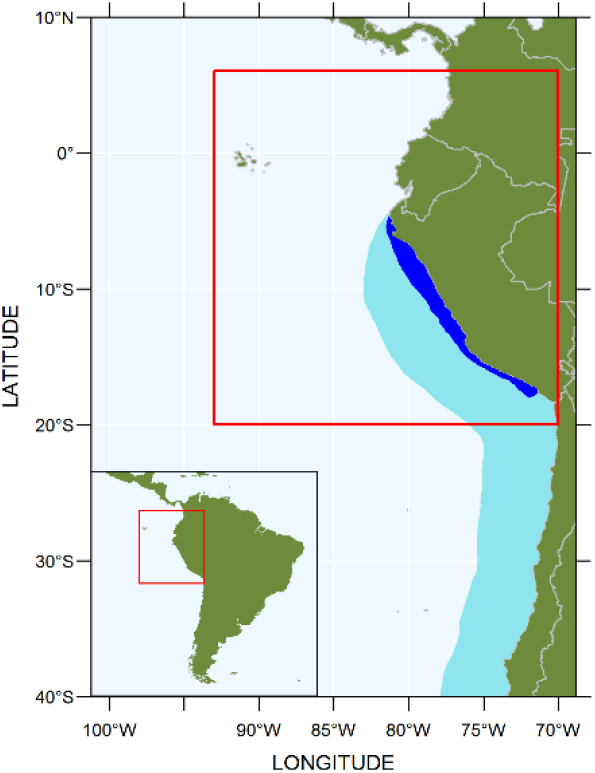
Map of the modeled area. The model spatial domain is limited by the red square. The light blue area shows the extension of the Humboldt Current Large Marine Ecosystem, and the dark blue area the extension of the Peruvian Upwelling Ecosystem.

**Figure 2:**
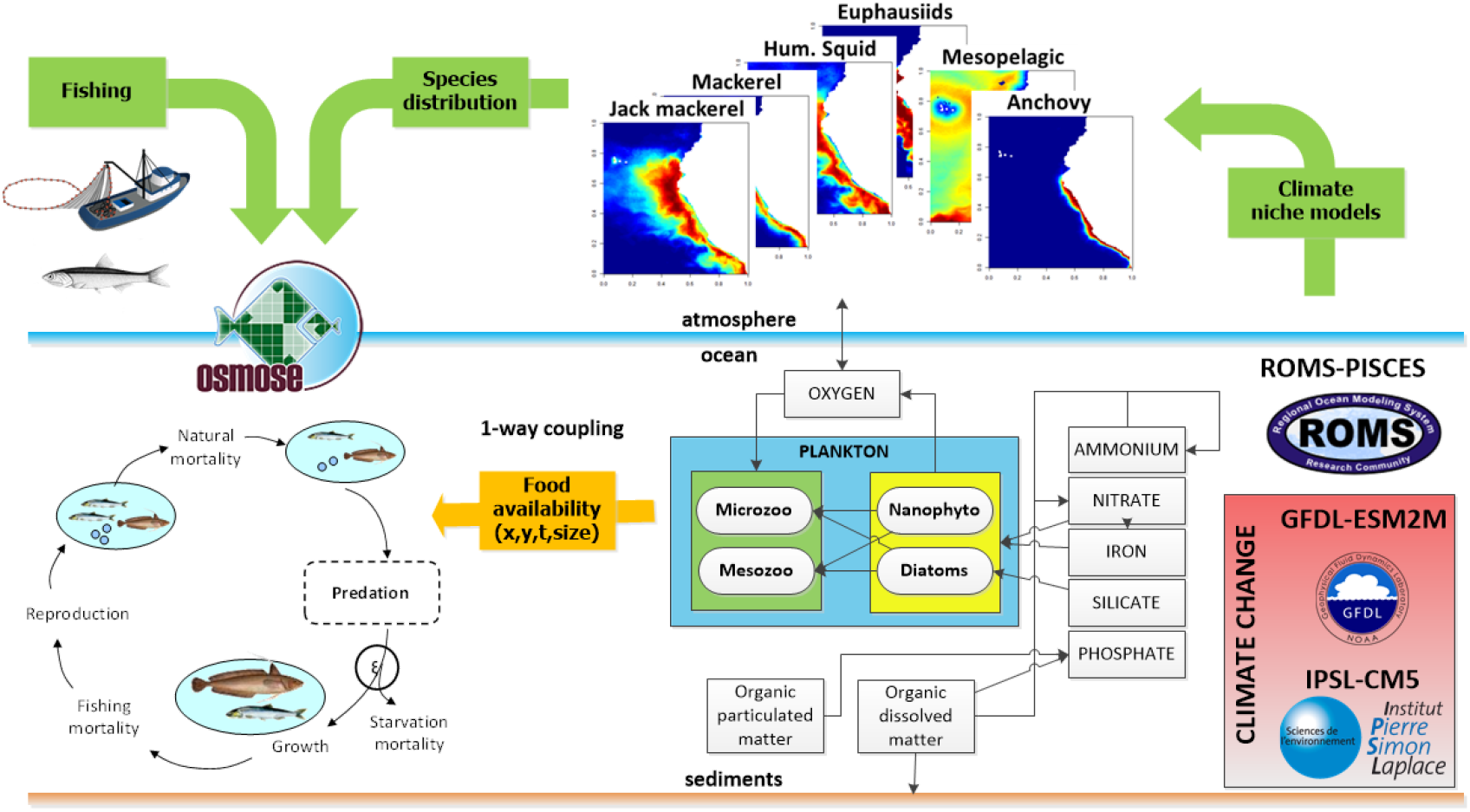
Conceptual representation of the modelling approach for projecting the impacts of climate change scenarios on Peruvian anchovy. The ESMs and the ROMS-PISCES model are used to provide plankton fields for OSMOSE and environmental inputs for the Climate Niche Models. Plankton biomass, species distribution maps and fishing are the main forcings of OSMOSE.

### The OSMOSE model

The ecosystem model OSMOSE (Shin and Cury 2001, 2004) is an individual based model for the simulation of exploited marine ecosystems. The basic unit of simulation is the “school”, a group of individuals of the same species sharing the same properties and history in terms of spatial position, length and age. The state of each school in the system can be described by a vector S = (s,x,y,N,L,A), where s is the species the school belongs to, (x,y) is the position of the school (longitude, latitude), N is the number of individuals in the school (abundance), L is the body length of the individuals and A is its age (Oliveros–Ramos 2014). At any time, the state of the system can be described by the state of all living schools.

There are three main processes controlling the dynamic in the model: mortality, growth and movement. The core of the first two processes relies in very simple survival and growth assumptions: exponential survival and von Bertalanffy growth (Shin and Cury 2001, 2004). These mortality and growth processes are formulated so they both depend on the predation process (predation success) which is length based in OSMOSE. For each species, a school can feed on prey within a limited range of sizes, parameterized by the minimum and maximum ratio between predator and prey sizes. This rule allows each school to select which other schools it can feed on, considering only the length of the individuals and the co-occurrence in the same cell, so that for most modeled species in the pelagic column, no a priori species-specific trophic relationship is assumed. As a result, the diets, trophic levels and predation mortalities are derived quantities from the model and the size-based predation assumptions and, produced as outputs of OSMOSE. The total mortality for a school is calculated taking into account the extraction from co-occurring predator schools (predation mortality), the fishing mortality, the starvation mortality and an additional mortality component M_0_ representing the mortality due to other processes which are not fully explicit in the model (e.g. due to other predators). The total mortality and its components for each school in the same cell of the grid are solved simultaneously for all species. In OSMOSE, the total mortality and growth in length during a time step depends on the state of the school itself and that of all other schools, regardless of the species, in a defined neighborhood (square), given by the grid of the spatial domain. This means growth and mortality are function of the state of all schools which are present in the same cell of the grid at the same time. In the OSMOSE version considered in this work (version 3.2), the functions defining mortality and growth are deterministic, while the main source of stochasticity is in the movement process (Oliveros–Ramos 2014). Additionally, the model is forced by species-specific spatial distribution maps. At any time, each school can be assigned to a unique map, while this map can change during the simulation according to an age and time-specific criterion (Oliveros–Ramos 2014). When a change in the map for a school occurs, the school is relocated randomly in the new map according to its spatial probability distribution. An additional important process leading to the renewal of the population is the reproduction process. Here, the total spawner biomass of each species (aggregated over all schools given a size or age of maturity and sex ratio) leads to an egg production (age 0 abundance). While the recruitment level emerges from the different sources of mortality applied at the subsequent time steps, the initial total egg production is assumed proportional to the spawner biomass, the fecundity potential (number of eggs per gram of mature female by unit of time) being the factor of proportionality. These eggs are distributed in a number of new schools proportional to the area of distribution of the species at age 0 and the expected average biomass of the population for the modeled period. This means that for species with bigger distribution areas, abundance, or both, more new age-0 schools are introduced in the model at each time step. The initial state of any new school is determined by its: (i) abundance: the number of eggs after the distribution of the total egg production among all the new schools, (ii) length: the assumed size of the egg for the species, (iii) position: randomly distributed within the age-0 map for the species, and (iv) age 0 (Oliveros–Ramos 2014). The dynamics of the low trophic level (LTL) species is not explicit in the model, reason why plankton fields (provided by observations or biogeochemical models) are used as forcing variables, representing additional food for the smaller species or size classes in OSMOSE. For the hindcast simulation (base model) we used the ROMS-PISCES model (Espinoza et al. 2016) and for the future forecast the Earth system models described in the next section.

The modeled area ranges from 20ºS to 6ºN and 93ºW to 70ºW covering the extension of the Northern Peru Current Ecosystem and the Peruvian Upwelling Ecosystem, with 1/6º of spatial resolution (Oliveros-Ramos et al. 2017). The model explicitly model the life history and spatio-temporal dynamics of 9 species (1 macro-zooplankton group, 1 crustacean, 1 cephalopod and 6 fish species: anchovy, sardine, jack mackerel, chub mackerel, hake and mesopelagic fishes), between 1992 and 2008.

### Earth system models

Earth system models (ESM) are the main tools to forecast the impact of Climate Change on Earth. Several ESM have been developed, but not all of them simulate the input necessary to force ecosystem models (Tittensor et al. 2018). For this reason, the models IPSL-CM5A-LR (??) and GFDL-ESM2M were used, as they provide all the inputs needed to force the OSMOSE model. The IPSL-CM5A-LR features a relatively strong surface warming and global NPP decline in future, while GFDL-ESM2M projects relatively small changes in both variables globally (Tittensor et al. 2018). Each model has been run using different scenarios of greenhouse gas concentrations (GHG) and radiative forcing, resulting in future climate changes, and ranging from the low emissions Representative Concentration Pathway RCP 2.6 scenario to the high emissions RCP 8.5 scenario. The description of the four RCP scenarios considered is shown in Table 2. One of the main limitations for the use of ESMs to force regional fish and fishery models is the coarse resolution at which the outputs are produced, 1ºx1º for the models considered here (ISIMIP 2a outputs), while missing important coastal areas where main fisheries are active.

**Table 2:** Descriptions of the climate change scenarios used (from GFDL and IPSL models).

| Scenario | Description | Reference |
| --- | --- | --- |
| RCP 2.6 (low emissions) | Scenario where GHG emissions are reduced over time, reaching a peak of 3 W/m<sup>2</sup> of radiative forcing by mid 21th century and stabilizing to 2.6 W/m<sup>2</sup> by 2100. | Van Vuuren et al. 2011 |
| RCP 4.5 (medium-low emissions) | Scenario where GHG emissions are reduced by different strategies and technologies to stabilize at 4.5 W/m<sup>2</sup> of radiative forcing by 2100. | Thomson et al. 2011 |
| RCP 6.0 (medium emissions) | Scenario where GHG emissions are reduced by different strategies and technologies to stabilize at 6.0 W/m<sup>2</sup> of radiative forcing after 2100 without exceeding the 6.0 W/m<sup>2</sup> before 2100. | Masui et al. 2011 |
| RCP 8.5 (high emissions) | Scenario where GHG gas emissions are increased over time, leading to high emission levels and reaching 8.5 W/m<sup>2</sup> by 2100. | Riahi et al. 2011 |

Additionally, since ESMs have not been specifically designed to force ecosystem models, some outputs such as planktonic biomass may have spurious or unrealistic values and needed to be checked before use (Tittensor et al. 2018). Taking into account these issues, we used the bias correction and statistical downscaling (BCSD) procedure described in Oliveros-Ramos et al. (submitted) in order to: i) increase the resolution of the ESM inputs, ii) reproduce the patterns of the coastal areas and, iii) correct bias in the ESM inputs for our model area. For the current work, we built a BCSD model for seven variables: the four plankton groups from the ESM (small and large groups of phyto and zooplankton), sea surface temperature (SST), integrated net primary productivity (NPP) and sea surface salinity (SSS). The high-resolution variables used to build the model at the target higher resolution (Table 3) were: ROMS-PISCES plankton groups, Reynolds Optimal Interpolation for the SST, SODA reanalysis for the SSS and the VGPM algorithm for the NPP (Ocean Productivity 2017). The BCSD was based on Generalized Additive Models (GAMs) using the mgcv approach (Wood 2011) implemented in R (R Core Team 2017). The model included spatially explicit covariates (latitude, longitude, distance to the continental shelf) and the main variables modeled: Sea Surface Temperature (SST), Sea Surface Salinity (SSS), Net Primary Production (NPP), Phytoplankton (small – SPHY, large LPHY) and zooplankton (small – SZOO, large LZOO).

**Table 3:** Environmental data inputs used for the bias correction and statistical downscaling models.

| Input variable | Source | Reference |
| --- | --- | --- |
| Sea Surface Temperature | OISST 0.25°x0.25° resolution, monthly averages. | Banzon et al. 2016, Reynolds et al. 2007 |
| Sea Surface Salinity | SODA 3.3.1 0.5°x0.5° resolution, monthly averages. | Carton and Giese 2008 |
| Net Primary Production | VGPM 1/6°x1/6° resolution. | Ocean Productivity 2017 |
| Phytoplankton | ROMS-PISCES 1/6°x1/6° resolution, monthly averages. | Espinoza et al. 2016 |
| Zooplankton | ROMS-PISCES 1/6°x1/6° resolution, monthly averages. | Espinoza et al. 2016 |

The BCSD models were used to do the forecast and produce the high-resolution inputs for the future emissions scenarios (RCP 2.6, 4.5, 6.0 and 8.5).

### Spatial distribution of fish

We used the statistical niche models for several key species of the Peruvian Upwelling Ecosystem developed by Oliveros-Ramos (2014). The models were constructed by logistic regression using GAMs. The dependent variable was the occurrence of the species in the acoustic surveys of IMARPE (Peruvian Marine Research Institute), and the predictors the monthly averages of remote-sensing and reanalysis environmental data (SST, SSS and NPP, see Table 3). The best model for each species was selected using the AUC in the validation set (random 30-70 split for training/validation) as detailed in Oliveros-Ramos (2014). These models were used to predict the spatial distribution in the forecast period using the inputs taken from the ESMs, after the same bias correction and downscaling process. We built maps for the spatial distribution of the species explicitly modeled in OSMOSE. Providing the probability of occurrence of a species given some environmental predictors (temperature, salinity, primary productivity and bathymetry), annual maps were produced with seasonal time resolution for all species (4 maps per year) except euphausiids, for which monthly resolution (12 maps per year) was used for the historical period because of their short lifespan and high turnover rate. For the future period, maps were generated with yearly time resolution for all species (1 map per year) except euphausiids, for which seasonal resolution (4 maps per year) was used. The probabilities predicted by the anchovy environmental niche model were used to estimate the Habitat Suitability Index (HSI), using the formula:

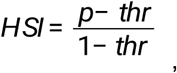

where *thr* (0.63) is the probability threshold which maximizes the skill of the model considering the area under the ROC curve (AUC) criterion for anchovy (Oliveros-Ramos 2014).

### Impact of environmental variability on anchovy population dynamics

One of the main drivers of the anchovy population dynamics is the larval mortality, controlling the level of production of new fish to the system (recruitment). For the historical period, the larval mortality was estimated as a set of time-varying parameters (Oliveros-Ramos et al. 2017) but in order to build a forecast model we modelled the time variability of the mortality(?) as a function of environmental variables. An index for the selected environmental variables (SST, NPP and SSS) was calculated as the average inside and outside the continental shelf (up to 200m of depth) to be used as independent variables to explain the larval mortality. For the more coastal species (anchovy, hake, munida, sardine and euphausiids) the inside-shelf index was used, while the outside-shelf index was used for the remaining species (horse mackerel, jack mackerel, mesopelagics and giant squid). These indices were calculated for each climate model (IPSL and GFDL) for the period 1992-2005, corresponding to the historical period of the simulations. We fitted one GAM model for each ESM model (IPSL and GFDL) including all variables but performing automatic ridge selection so the non-significant variables are weighted out of the model (by taking their coefficients close to zero), fitting the following model:

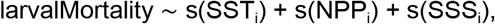

where the *i* index can take the values for inside or outside the shelf depending on the modeled species. For the forecast, the results of the statistical downscaling were used to compute the same index over the continental shelf (up to 200 m of depth) and outside this area and used it to predict the larval mortality for the future scenarios.

### Fishing

We used a simple *business as usual* (BAU) approach for fishing, since fisheries management regulations are complex, especially during low abundance periods. We used the average of 2005-2008 for the fishing mortality for every species in the model. For anchovy, this 2005-2008 average represented a conservative value.

## Results and discussion

The anchovy’s habitat showed a reduction in quality (HSI) and area (AREA) for all emissions scenarios combined with fishing BAU (Figure 3), from minor changes using the Earth system model GFDL to more than 30% using the IPSL climate model under the RCP 8.5 scenario, with major changes observed after 2050. Both habitat variables showed a decreasing linear trend with the relative change of sea surface temperature (SST), at a rate of −19.6% for the relative change in the HSI and −17.6% for the relative change in the area of the habitat. They increased in a non-linear way with the relative change in net primary productivity (NPP), while they remained almost constant over a large range of NPP decrease and only started to decrease when the decrease of NPP was over 35% of the current values. The net secondary production (NSP) of anchovy showed a higher uncertainty than the spatial distribution, including extinction of the population for the more drastic changes in temperature, with a significant linear trend of −65% per relative change in SST, but with zero production over the 50% of increase in temperature (equivalent to an average temperature over the shelf of around 32ºC). However, the collapse in the secondary production can be observed for smaller, more likely, changes in temperature, related to the cumulative impacts of smaller warming conditions in the population dynamics of anchovy, as observed mainly for the high GHG emissions RCP 8.5 scenario. An increase in NSP is expected with the increase in NPP. For both explanatory variables (SST and NPP) a high uncertainty in the relationship is observed (Figure 3). These higher levels of uncertainty with respect to the NSP can be related to the impact of trophic interactions which also affect anchovy productivity.

**Figure 3.**
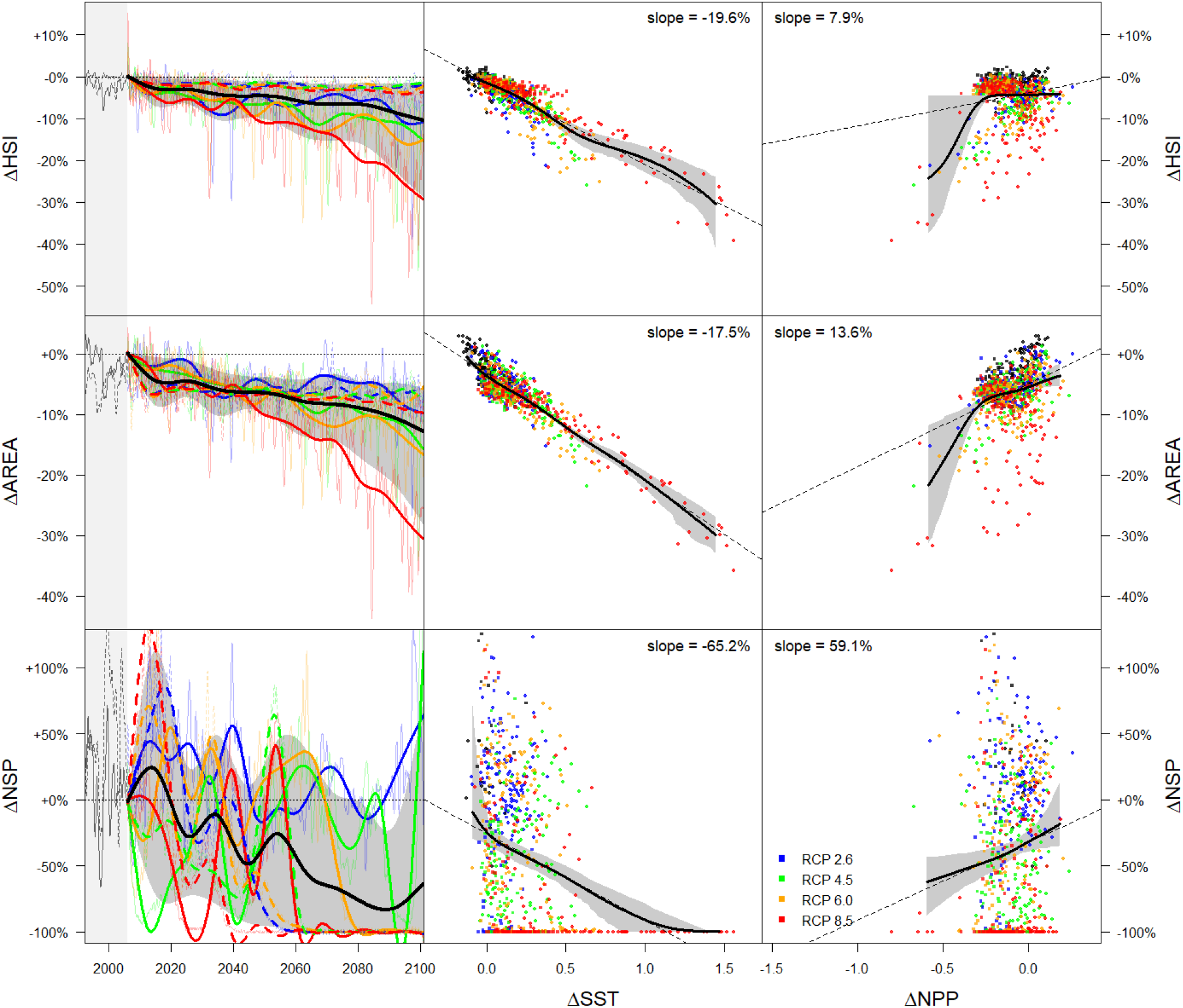
Impact of climate change scenarios on the distribution and productivity of Peruvian anchovy. We analyzed the relative change in the habitat suitability index (HSI, first row, a-c), the extent of anchovy occurrence (second row, d-f) and anchovy’s net secondary production (NSP, third row, g-i). The first column shows the temporal variability of the mentioned variables for all the climate models and scenarios (solid line: IPSL, dotted line: GFDL). The second and third columns show the relative change of the variables given the relative change of temperature and net primary production, respectively, for all the climate models and RCP scenarios.

The changes in the spatial distribution of anchovy are also seen as changes in the spatial distribution of landings, as we can observe in Figure 4, with gravity centers of landings showing a shift to the south and more coastal areas for all future GHG emissions scenarios, while more variability is observed in the results forced by the IPSL model. On average, a small displacement to the south and to more coastal areas is expected. These may have an economic impact on the fishery, since latitudinal distance of the trips would be expected to increase, since major fishery ports and processing plants have fixed positions in the short term. Additionally, a more coastal distribution can increase the mixing between juveniles and adults, particularly in warming El Niño-like conditions and increase the depth of the schools (Ñiquen and Bouchon 2004), complicating the fisheries activity and potentially increasing by-catch of anchovy juveniles.

**Figure 4.**
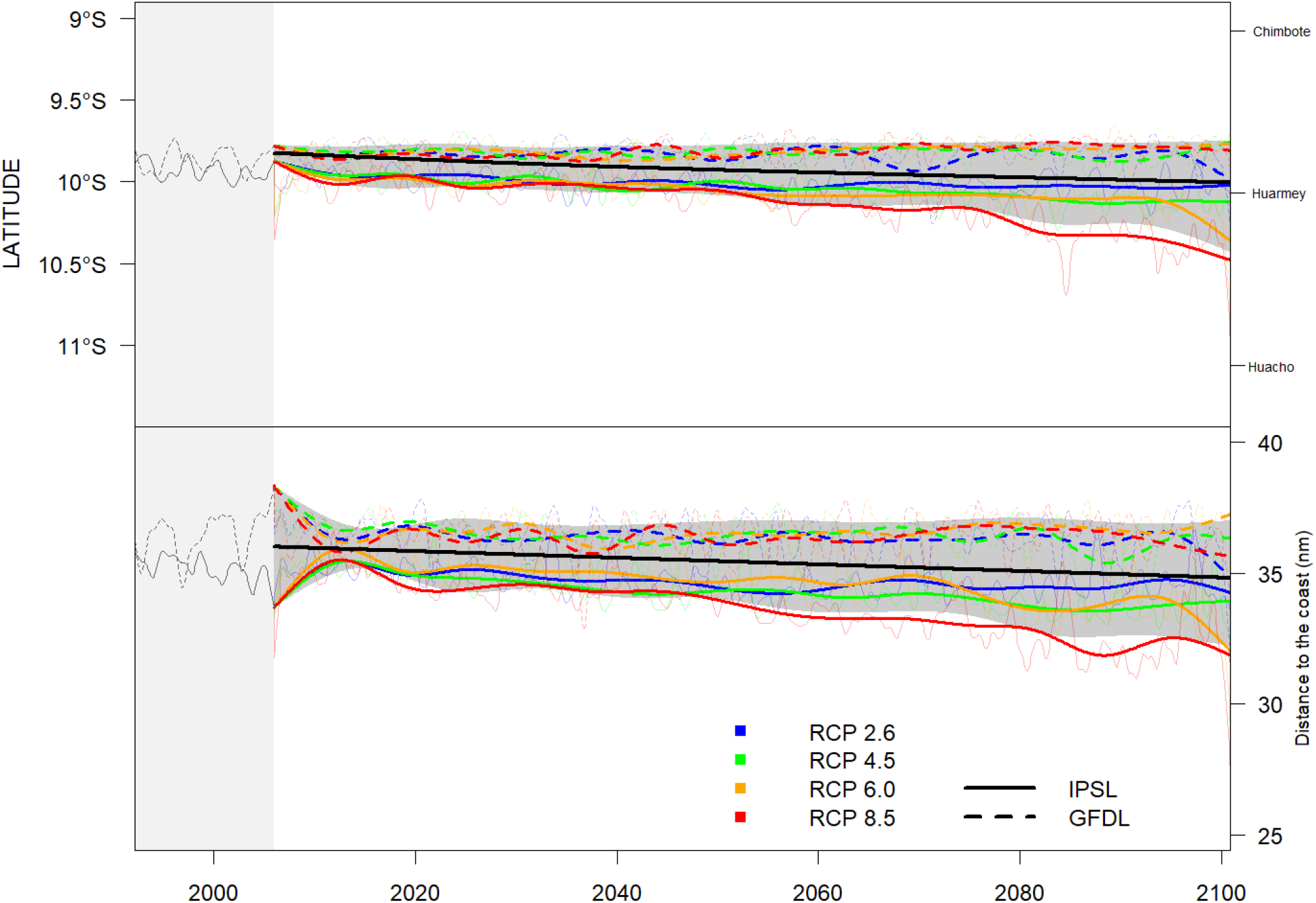
Gravity center of anchovy landings under RCP scenarios and Fishing BAU. The gravity center in terms of latitude (top) and distance to the coast (bottom) is shown for all the climate models (IPSL solid line, GFDL dotted line) and RCP scenarios.

**Figure 5.**
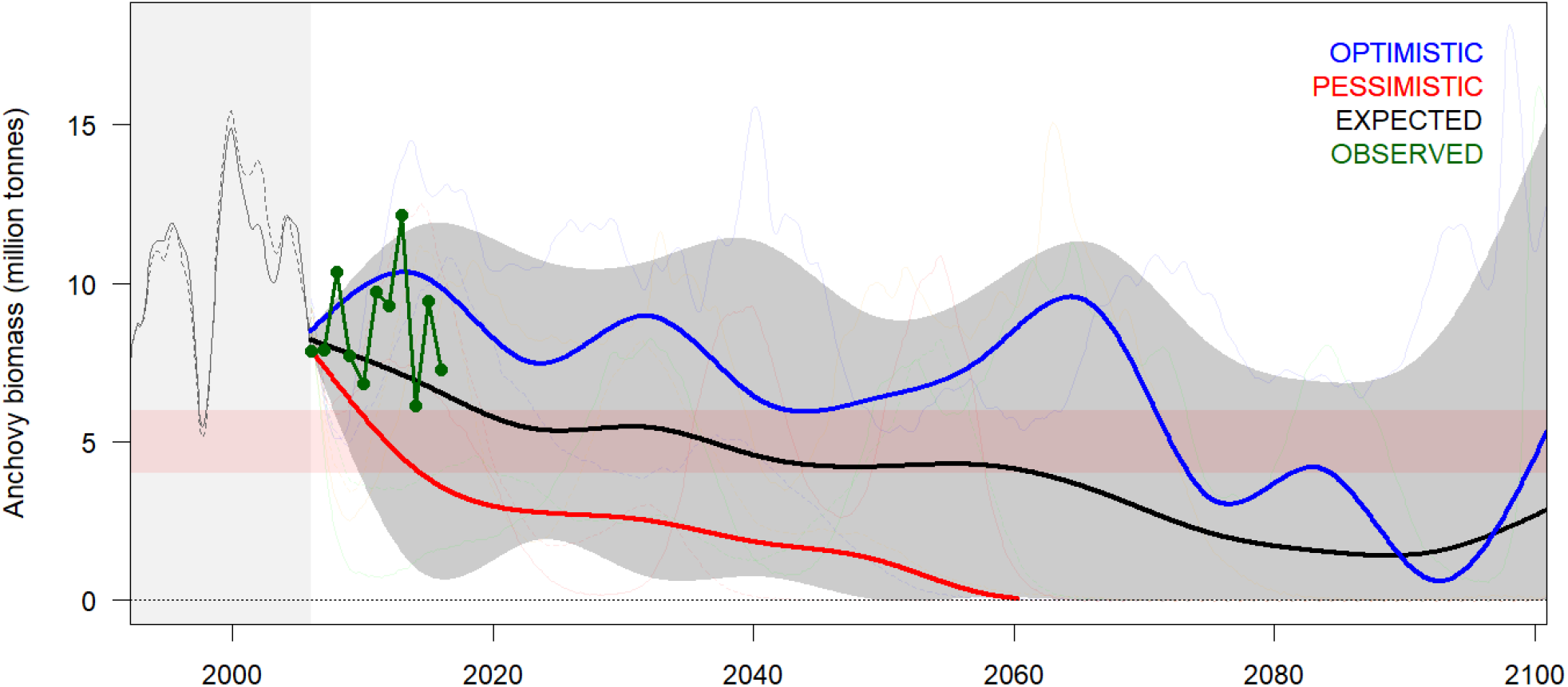
Impact of RCP scenarios in the relative change of Peruvian anchovy biomass. The lines represent the mean (black), 20% quantile (red) and 80% quantile (blue) of all simulations, while the shaded area is the 95% envelope of all simulations. The 20% quantile is considered the pessimistic trajectory and the 80% quantile the optimistic trajectory, given all the scenarios considered. Observed biomass of anchovy from acoustic surveys for the recent years (2006-2016) is shown in green.

The simulations also showed that anchovy biomass was projected to eventually collapse (by dropping below the biomass limit reference point of 4 to 6 millions tonnes) for most scenarios. By taking the lower 20 percentile of biomass distribution (grouping all anchovy biomass produced by the RCPs and climate models and OSMOSE replicated runs) as pessimistic and the upper 80 percentile as optimistic, our results showed a likely collapse of the anchovy population by 2020 and near extinction by 2060 in the pessimistic case. For the optimistic case, the population does not collapse before 2070; while on average during the 2020 to 2060 period the population remains sustainable, above the biomass limit reference point. The simulations also predict a potential recovery by the end of the century, related to the favorable conditions expected under the RCP 2.6 scenario, where the greenhouse gas emissions are reduced over time and stabilized after mid-21st century. Considering that between 2006 to 2016 the anchovy biomass has been between the recent range of variability and has been estimated to an average level of 8.1 million tonnes (IMARPE 2017) the current status seems to be between the average and optimistic trajectories. However, it is important to consider that our simulations used a *business as usual* scenario for fishing which strongly differs from the adaptative fisheries management implemented by IMARPE and the Peruvian Government to set the total allowable catch every year (IMARPE 2016), especially reductions of catch implementations during El Niño events. In this regard, the impact of adaptive management to ensure the sustainability of exploited resources under climate change scenarios should be explored in future scenarios studies.

The optimistic trajectory of anchovy biomass represents a reduction at a rate of 14% per decade until mid 21st century, while no changes in the spatial distribution of the population or landings were observed. In this optimistic trajectory, adaptive fishery management could have options to increase the sustainability of the resource. In a pessimistic scenario, the anchovy biomass reduces at a rate of 22% per decade, with a collapse after 2020 and near extinction by 2060. A similar trend with biomass below the biological reference point has been observed historically during the warmer regime of the 1980s, followed by a recovery by mid 1990s. However, this recovery followed a near fishing moratorium for anchovy (with a shift to sardine exploitation) and a cooling of the system leading to a new colder regime after El Niño 1997-1998 (Ñiquen and Bouchon 2004). So, while a recovery is theoretically possible, an increasing impact of anthropogenic climate change is not compatible with such recovery situation while management procedures applied can be very important to ensure the sustainability of the stock. The latter shows an important limitation of this study, as the *business as usual* scenario for fishing cannot represent the diverse array of possible fisheries responses under different climate and anchovy biomass scenarios, particularly for a near-collapse situation. Another limitation of this study is to have ignored oxygen as a driver of anchovy dynamics due to the lack of future scenarios producing this key environmental variable (Bertrand et al. 2011). Also, a longer hindcast model simulation could be useful to better understand the environmental relationships between anchovy distribution and mortality, which are very important to better forecast the future of the population. This represents the first study on the climate change impacts on Peruvian anchovy population and highlights the requirements for the coupling of Earth system models with regional models in order to produce realistic projections of future impacts of global change on key resources such as the Peruvian anchovy.

### Conclusions

Our results suggested climate change is expected to have a negative impact on Peruvian anchovy under all GHG emissions RCP scenarios combined with BAU fishing, while the strength of the impact has a high uncertainty; from an optimistic trajectory where a reduction in the anchovy biomass is expected under *business as usual* fishing conditions followed by a recovery after mid-21st century; to a pessimistic trajectory where anchovy fishery collapses within a decade, followed by a near-extinction of anchovy population off Peru. A more complex analysis of the impacts of fishing should be conducted to fully analyze the fishing impacts on the anchovy population, and several fisheries management procedures should be included in scenarios, in particular adaptive fisheries management strategy, in order to evaluate potential mitigation management solutions for the sustainability of anchovy population and the fisheries it supports.

## Acknowledgements

We thank the financial support from the IMARPE-PRODUCE-IADB Project “Adaptation to climate change of the fishery sector and marine-coastal ecosystem of Perú” (PE-G1001/PE-T1297). This work has been partially funded by the BiodivErsA and Belmont Forum project SOMBEE (BiodivScen programme, ANR contract n°ANR-18-EBI4-0003-01). We acknowledge the support of the Instituto del Mar del Peru (IMARPE) and the Institut de Recherche pour le Développement (IRD), through the LMI DISCOH, LMI ICEMASA and UMR MARBEC. We acknowledge the support of the Fish-MIP group.

## References

Aumont O., Bopp L., 2006. Globalizing results from ocean in situ iron fertilization studies, Global Biogeochem Cycles, 20, GB2017.

Barange, M., Merino, G., Blanchard, J. L., Scholtens, J., Harle, J., Allison, E. H., Allen, J. I., Holt, J. and Jennings, S. 2014. Impacts of climate change on marine ecosystem production in societies dependent on fisheries, Nat. Clim. Chang., 4(3), 211–216.

Bernhardt, J. R. and Leslie, H. M. 2013. Resilience to Climate Change in Coastal Marine Ecosystems, Ann. Rev. Mar. Sci., 5(1), 371–392, doi:10.1146/annurev-marine-121211-172411.

Bertrand A, Chaigneau A, Peraltilla S, Ledesma J, Graco M, et al. (2011) Oxygen: A Fundamental Property Regulating Pelagic Ecosystem Structure in the Coastal Southeastern Tropical Pacific. PLOS ONE 7(2): 10.1371

Blanchard, J. L., Jennings, S., Holmes, R., Harle, J., Merino, G., Allen, J. I., Holt, J., Dulvy, N. K. and Barange, M. 2012. Potential consequences of climate change for primary production and fish production in large marine ecosystems, Philos. Trans. R. Soc. B Biol. Sci., 367(1605), 2979–2989.

Chavez F.P., A. Bertrand, R. Guevara-Carrasco, P. Soler and J. Csirke. 2008. The northern Humboldt Current System: Brief history, present status and a view towards the future. Progress in Oceanography 79:95–105.

Cheung, W. W. L., Frolicher, T. L., Asch, R. G., Jones, M. C., Pinsky, M. L., Reygondeau, G., Rodgers, K. B., Rykaczewski, R. R., Sarmiento, J. L., Stock, C. and Watson, J. R. 2016. Building confidence in projections of the responses of living marine resources to climate change, ICES J. Mar. Sci., doi:10.1093/icesjms/fsv250.

Dufresne, J.-L., et al. 2013. Climate change projections using the IPSL-CM5 Earth System Model: from CMIP3 to CMIP5. Climate Dynamics, 40, 9-10, 2123-2165, 2013.

Dunne J.P et al. 2012. GFDL’s ESM2 Global Coupled Climate–Carbon Earth System Models. Part I: Physical Formulation and Baseline Simulation Characteristics. Journal of Climate, 25, 6646–6665.

Dunne J.P et al. 2013. GFDL’s ESM2 Global Coupled Climate–Carbon Earth System Models. Part II: Carbon System Formulation and Baseline Simulation Characteristics. Journal of Climate, 26, 2247–2267.

Echevin V., O. Aumont, J. Ledesma, G. Flores, 2008. The seasonal cycle of surface chlorophyll in the Peruvian upwelling system: A modelling study. Progress in Oceanography 79 (2008) 167–176.

FAO. 2016. The State of World Fisheries and Aquaculture 2016, Rome.

Halpern BS, Walbridge S, Selkoe KA, Kappel CV, Micheli F, et al. 2008. A global map of human impact on marine ecosystems. Science 319:948–52.

Hoar and Nychka. 2008. Statistical downscaling of the Community Clima System Model (CCSM) monthly temperature and precipitacion projections.

IMARPE. 2016. Elaboración de la Tabla de Decisión para la determinación del Límite Máximo de Captura Total Permisible para la pesquería del Stock Norte-Centro de la anchoveta peruana. IMP-DGIRP/AFDPERP edición 3v1.

IMARPE 2017. Análisis poblacional de la pesquería de anchoveta en el Ecosistema Marino Peruano. Informe del Instituto del Mar del Perú, http://www.imarpe.pe/imarpe/archivos/informes/info_anal_pob_anchov_1.pdf.

Masui et al. 2011. An emission pathway for stabilization at 6 $Wm^{-2}$ radiative forcing. Climatic change 109:59.

Lehodey, P., Senina, I., Nicol, S. and Hampton, J. 2015. Modelling the impact of climate change on South Pacific albacore tuna, Deep Sea Res. Part II Top. Stud. Oceanogr., 113, 246–259.

Mignot, J., et al. 2013. On the evolution of the oceanic component of the IPSL climate models from CMIP3 to CMIP5: A mean state comparison. Ocean Modelling, 72, 167-184, 2013, doi:10.1016/j.ocemod.2013.09.001

Mullon, C., Steinmetz, F., Merino, G., Fernandes, J. A., Cheung, W. W. L., Butenschen, M. and Barange, M. 2016. Quantitative pathways for Northeast Atlantic fisheries based on climate, ecological-economic and governance modelling scenarios, Ecol. Modell., 320, 273–291.

Oliveros-Ramos R. 2014. End-to-end modelling for an ecosystem approach to fisheries in the Northern Humboldt Current Ecosystem. PhD thesis, University of Montpellier 2, France. DOI: 10.13140/RG.2.2.34207.36001

Oliveros-Ramos R., Verley P., Echevin V., Shin Y-J. 2017. A sequential approach to calibrate ecosystem models with multiple time series data. Progress in Oceanography 151 (2017) 227–244.

Oliveros-Ramos R. et al. Submitted. Asynchronous statistical downscaling and bias correction by non-parametric distribution mapping for climate change impact applications.

Pörtner, H.-O., Karl, D. M., Boyd, P. W., Cheung, W. W., Lluch-Cota, S. E., Nojiri, Y., Schmidt, D. N. and Zavialov, P. O. 2014. Ocean Systems, in Climate Change 2014: Impacts, Adaptation and Vulnerability. Part A: Global and Sectoral Aspects. Contribution of Working Group II to the Fifth Assessment Report of the Intergovernmental Panel on Climate Chagne, edited by K. F. Drinkwater and A. Polonsky, pp. 411–484, Cambridge University Press.

R Core Team. 2017. R: A language and environment for statistical computing. R Foundation for Statistical Computing, Vienna, Austria. URL https://www.R-project.org/.

Riahi et al. 2011. RCP 8.5—A scenario of comparatively high greenhouse gas emissions. Climatic change 109:33.

Shin Y.-J., Cury P., 2001. Exploring fish community dynamics through size-dependent trophic interactions using a spatialized individual-based model. Aquatic Living Resources, 14(2): 65–80.

Shin Y.-J., Cury P., 2004. Using an individual-based model of fish assemblages to study the response of size spectra to changes in fishing. Canadian Journal of Fisheries and Aquatic Sciences, 61: 414–431.

Sumaila, U. R., Cheung, W., Dyck, A., Gueye, K., Huang, L., Lam, V., Pauly, D., Srinivasan, T., Swartz, W., Watson, R. and Zeller, D. 2012. Benefits of Rebuilding Global Marine Fisheries Outweigh Costs, PLoS One, 7(7), e40542.

Tittensor D., et al. A protocol for the intercomparison of marine fishery and ecosystem models: Fish-MIP v1.0. Geophysical Model Development. In press.

Thomson et al. 2011. RCP4.5: a pathway for stabilization of radiative forcing by 2100. Climatic change 109:77.

Travers M., Shin, Y.-J., Jennings, S., Machu, E., Huggett, J.A., Field, J.G., Cury, P.M., 2009. Two-way coupling versus one-way forncing of plankton and fish models to predict ecosystem changes in the Benguela. Ecological modelling 220: 3089–3099.

Travers-Trolet M., Y.-J. Shin and J.G. Field., 2013. An end-to-end coupled model ROMS-N2P2Z2D2-OSMOSE of the southern Benguela foodweb: parameterisation, calibration and pattern-oriented validation, African Journal of Marine Science, 36:1, 11–29.

United Nations> (UN). 2016. The first global integrated marine assessment: World Ocean Assessment I, New York.

United Nations Environment Program (UNEP). 2006. Marine and coastal ecosystems and human well-being: synthesis: a synthesis report based on the findings of the Millennium Ecosystem Assessment. World Conservation Monitoring Centre (WCMC), 155pp.

Van Vuuren et al. 2011. RCP2.6: exploring the possibility to keep global mean temperature increase below 2ºC. Climatic change 109:95

Wood, S.N. 2011. Fast stable restricted maximum likelihood and marginal likelihood estimation of semiparametric generalized linear models. Journal of the Royal Statistical Society (B) 73(1):3–36.

